# An Evaluation of Phylogenetic Workflows in Viral Molecular Epidemiology

**DOI:** 10.1101/2020.11.24.396820

**Authors:** Colin Young, Sarah Meng, Niema Moshiri

## Abstract

The use of computational techniques to analyze viral sequence data and ultimately inform public health intervention has become increasingly common in the realm of epidemiology. These methods typically attempt to make epidemiological inferences based on multiple sequence alignments and phylogenies estimated from the raw sequence data. Like all estimation techniques, multiple sequence alignment and phylogenetic inference tools are error-prone, and the impacts of such imperfections on downstream epidemiological inferences are poorly understood. To address this, we executed multiple commonly-used workflows for conducting viral phylogenetic analyses on simulated viral sequence data modeling HIV, HCV, and Ebola, and we computed multiple methods of accuracy motivated by transmission clustering techniques. For multiple sequence alignment, MAFFT consistently outperformed MUSCLE and Clustal Omega in both accuracy and runtime. For phylogenetic inference, FastTree 2, IQ-TREE, RAxML-NG, and PhyML had similar topological accuracies, but branch lengths and pairwise distances were consistently most accurate in phylogenies inferred by RAxML-NG. However, FastTree 2 was orders of magnitude faster than the other tools, and when the other tools were used to optimize branch lengths along a fixed topology provided by FastTree 2 (i.e., no tree search), the resulting phylogenies had accuracies that were indistinguishable from their original counterparts, but with a fraction of the runtime. Our results indicate that an ideal workflow for viral phylogenetic inference is to (1) use MAFFT to perform MSA, (2) use FastTree 2 under the GTR model with discrete gamma-distributed site-rate heterogeneity to quickly obtain a reasonable tree topology, and (3) use RAxML-NG to optimize branch lengths along the fixed FastTree 2 topology.

## Introduction

In molecular epidemiology, phylogenetic analyses are performed on viral sequence data to help reconstruct the evolutionary history and patterns of transmission for highly contagious diseases, providing valuable insights which may inform intervention strategies (Hall 2013) A standard molecular epidemiology workflow typically consists of the following: (1) multiple sequence alignment (MSA), (2) phylogenetic inference, (3) rooting, (4) dating, and (5) transmission clustering. Since each step in the workflow occurs downstream of other analyses, the accuracy of each step is largely dependent on the accuracy of the steps preceding it. Notably, MSA and phylogenetic inference play an integral role in transmission clustering techniques utilized in molecular epidemiology. For example, HIV-TRACE is typically considered the best-practice method for HIV transmission clustering: pairwise distances are estimated from a MSA of viral genomes collected from individuals, and two individuals are considered to be epidemiologically “linked” if the pairwise distance between their viral samples is below a user-specified threshold (Kosakovsky *et al*. 2018). TreeCluster (Balaban *et al*. 2019), Cluster Picker (Ragonnet-Cronin *et al*. 2013), and PhyloPart (Prosperi *et al*. 2011) are transmission clustering tools that separate individuals based on the phylogenetic relationships between viral samples, typically relying on tree topology, branch lengths, and pairwise distances.

Both MSA and phylogenetic inference are NP-Hard computational problems (Chatzou *et al*. 2016), meaning optimal solutions are computationally infeasible on even relatively small datasets. As a result, several heuristic tools have been developed to provide near-optimal approximations. Notably, viral MSA is commonly conducted using MAFFT (Katoh and Standley 2013), MUSCLE (Edgar 2004), and Clustal Omega (Seivers *et al*. 2011), and viral phylogenetic inference is commonly conducted using IQ-TREE (Chernomor *et al*. 2016), FastTree 2 (Price *et al*. 2010), RAxML-NG (Kozlov *et al*. 2019), and PhyML (Guindon *et al*. 2019). As a result, researchers have multiple potential phylogenetic inference workflows at their disposal, each with a theoretical trade-off between runtime and accuracy, yet the impacts of inherent errors and imperfections in these heuristic estimation approaches on downstream molecular epidemiological results are poorly understood.

In this manuscript, we execute multiple commonly-used workflows for conducting viral phylogenetic analyses on simulated viral sequence data modeling HIV, HCV, and Ebola, and we compute multiple methods of accuracy motivated by transmission clustering techniques.

## Methods

Our simulated datasets were motivated by curated alignments of real world sequence data: from the Los Alamos National Laboratory (LANL), we obtained full genome MSAs of Ebolavirus (710 total), Hepatitis C Virus (HCV, 471 total), and Human Immunodeficiency Virus Type 1 (HIV-1, 4004 total).

From each curated MSA, we used IQ-TREE to infer a phylogeny under the General Time Reversible (GTR) model of sequence evolution (Tavaré 1986) with invariable sites and gamma-distributed site-rate heterogeneity with 20 categories. From the IQ-TREE results, we obtained the phylogeny, the GTR substitution model parameters, the proportion of invariable sites, and the shape of the gamma distribution. The phylogeny was subsequently rooted using minimum variance rooting (Mai *et al*. 2017), and a tree with 100 leaves was subsampled from it (Fig. 1). These parameters were then used to simulate sequence alignments from the subsample of the inferred phylogeny using INDELible (Fletcher & Yang 2009). 10 replicate datasets were generated for each virus.

**Figure 1.**
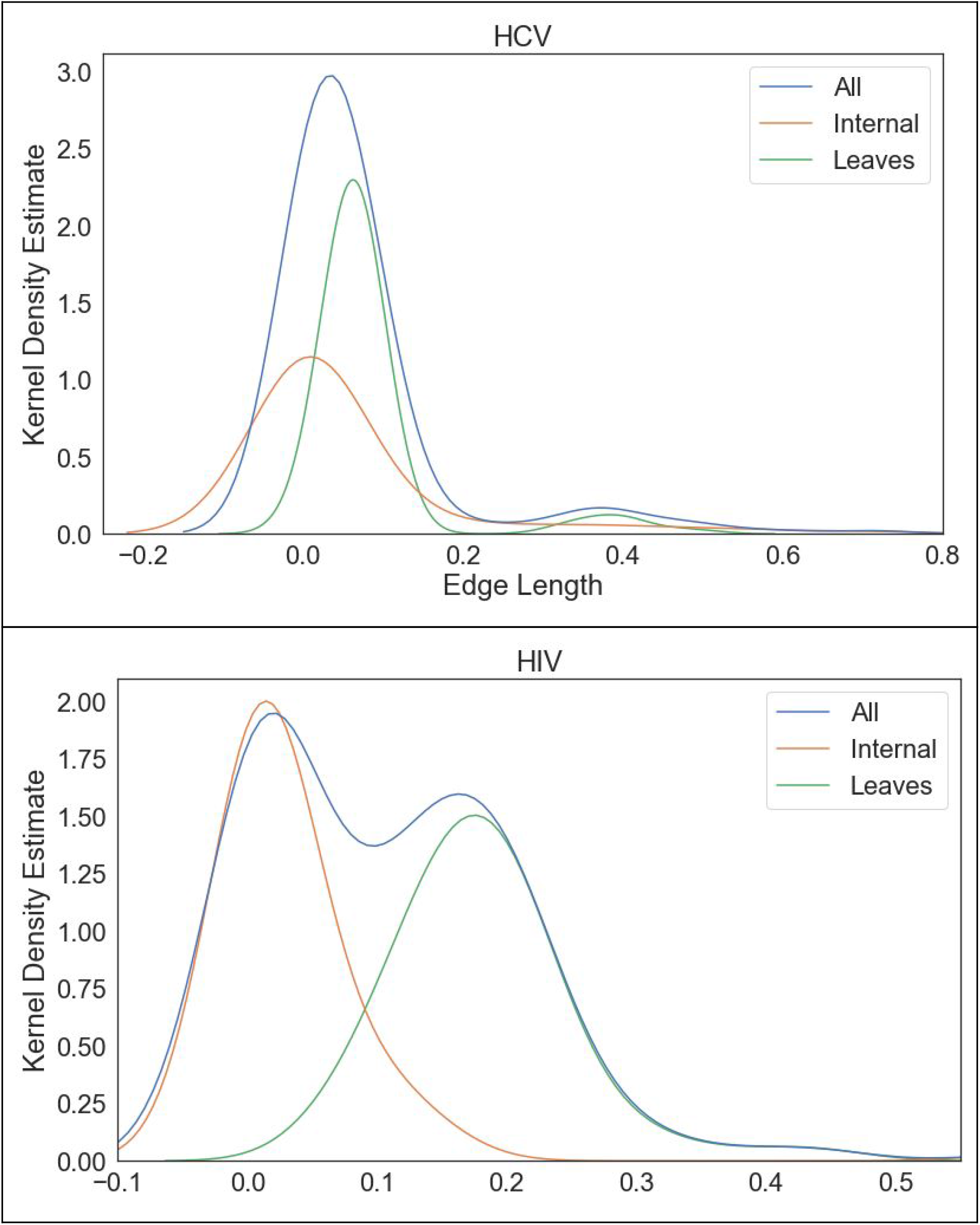
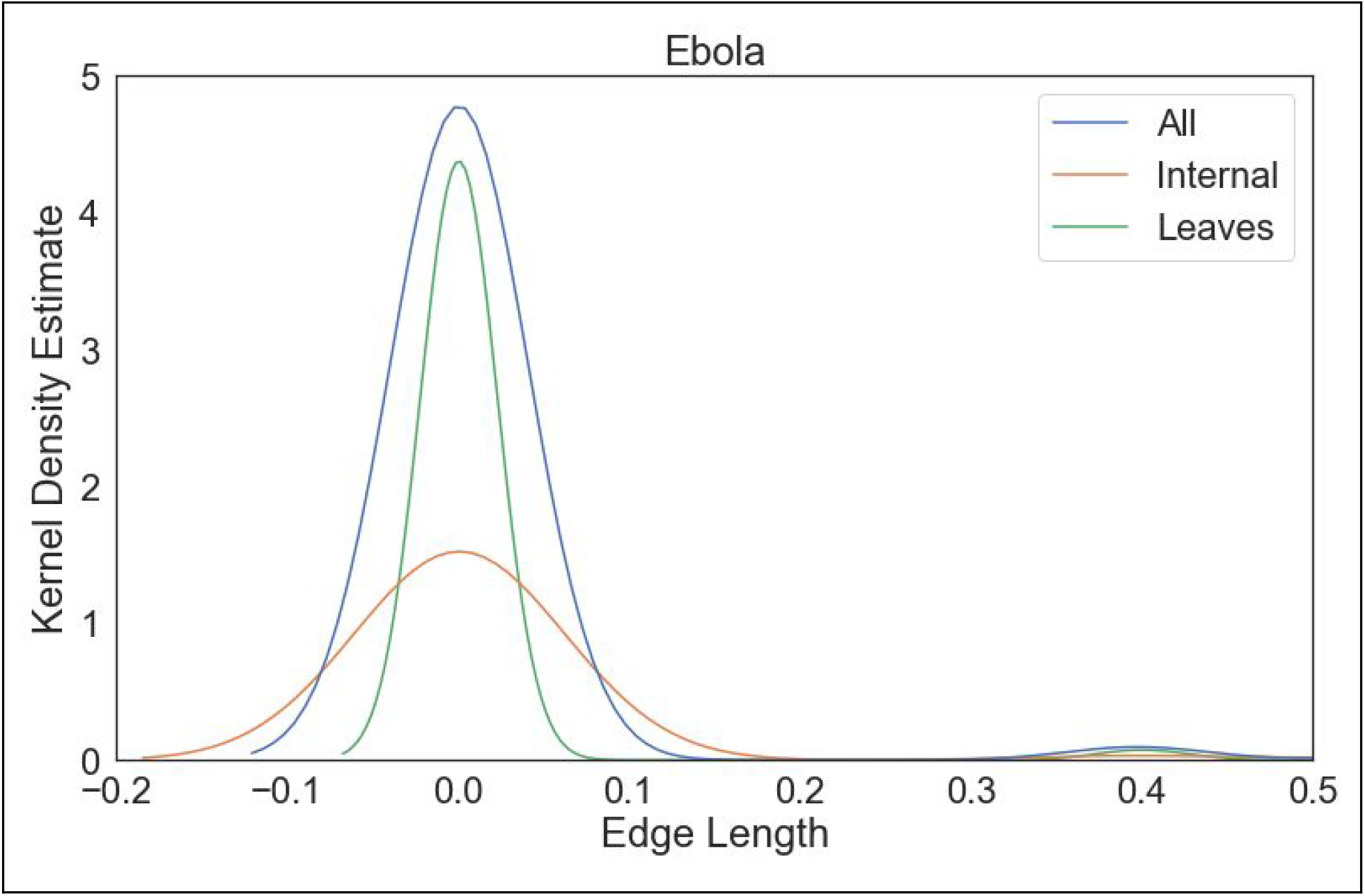
Kernel density estimates of the branch length distributions for the Ebola, HIV, and HCV true phylogenies.

**Figure 2.**
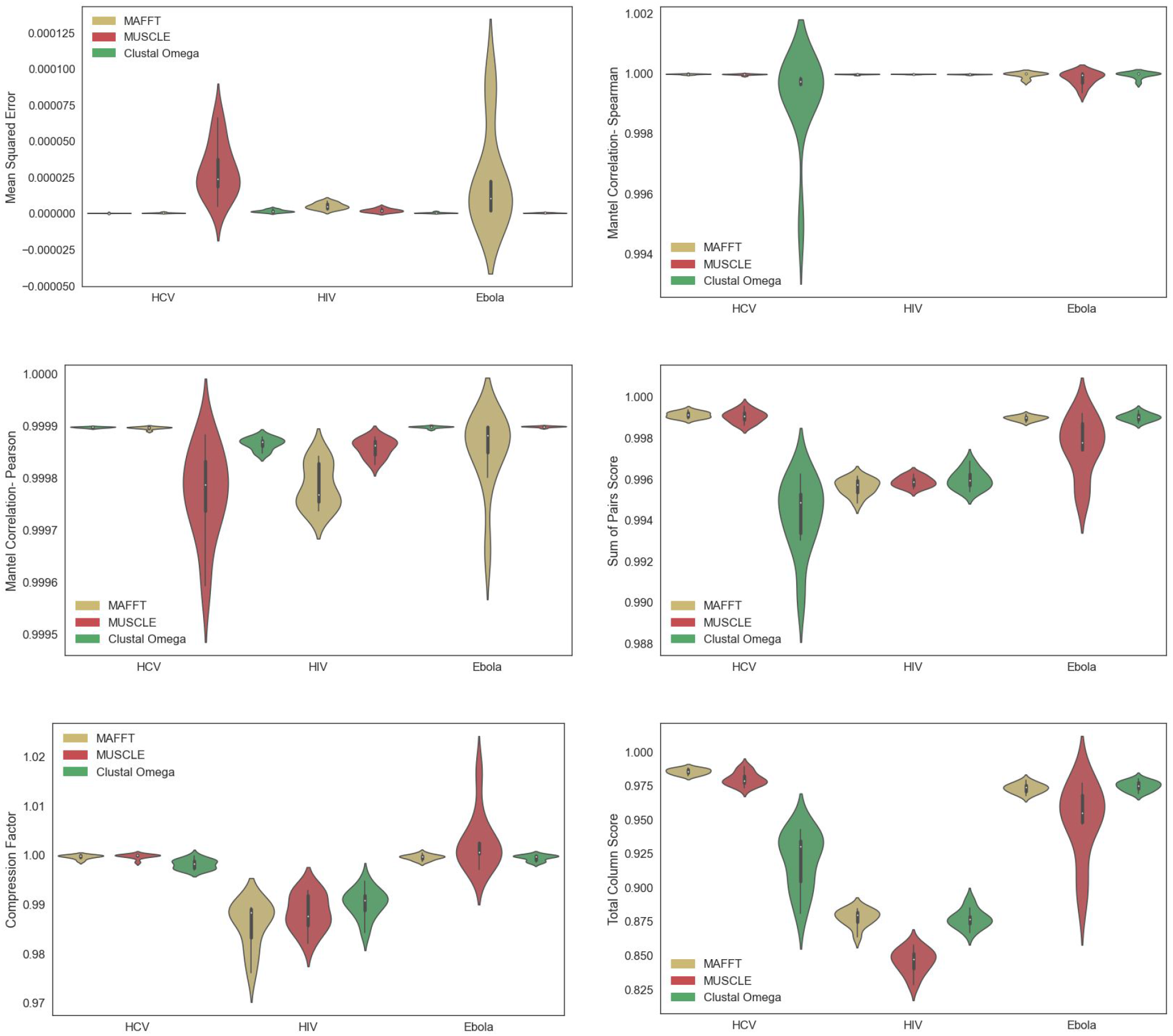
Metrics of sequence alignment accuracy for MAFFT, MUSCLE, and Clustal Omega on 10 simulated replicate datasets of HIV, HCV, and Ebola. Violin plots are shown for Mean Squared Error, Spearman/Pearson Mantel Correlation, SP score, TC score, and Compression Factor.

We then utilized the true simulated alignment produced by INDELible to evaluate the accuracy of the alignments estimated by MAFFT, Clustal Omega, and MUSCLE.

To assess the accuracy of phylogenetic inference tools, we ran FastTree 2, IQ-TREE, RAxML-NG, and PhyML on the true MSAs and compared the results against the tree along which the sequences were simulated. All tools utilized the GTR model of sequence evolution. FastTree 2 and PhyML utilized discrete gamma-distributed site-rate heterogeneity, whereas RAxML-NG and IQ-TREE utilized the FreeRate model (Yang 1995). We also utilized IQ-TREE’s ModelFinder Plus functionality (Kalyaanamoorthy *et al*. 2017).

In addition to the above analyses, in which we allowed each phylogenetic inference tool to perform tree search, we also examined a workflow in which a phylogeny is first inferred using FastTree 2, and then each of the other inference tools simply optimizes the branch lengths using the FastTree 2 phylogeny as a fixed tree topology (i.e., the second tool does not execute tree search). Given that FastTree 2 is an order of magnitude faster than the other tools (Tab. 1), this could potentially be an effective way to reduce computational time for phylogenetic inference workflows.

**Table 1.** Total runtime for phylogenetic inference (top row) and runtime of branch length optimization on a fixed topology (bottom row) for FastTree 2, RAxML, IQ-TREE (GTR), and IQ-TREE MFP on a curated MSA of 2,322 HIV-1 whole genome sequences from LANL. PhyML was unable to execute due to high memory consumption. All runs were executed sequentially on a 4-core 3.5 GHz Intel i5-6600k with 16 GB of memory, and each tool automatically selected an optimal number of threads to use internally.

| (Seconds) | FastTree 2 | RAXML | PhyML | IQ-TREE | IQ-TREE MFP |
| --- | --- | --- | --- | --- | --- |
| Total | 645 | >604,800 | memory | 84,931 | 266,399 (142,286 MFP) |
| BL optimization |  | 757 | memory | 1,532 | 4,885 |

Lastly, to assess the accuracy of different combinations of tools, we ran each phylogenetic inference method on the sequences aligned by each MSA tool and similarly measured the accuracy of each inferred phylogeny. The metrics used to assess accuracy are described below.

### Mean Squared Error

We use Mean Squared Error as a measure of patristic distance matrix similarity

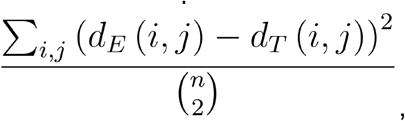

where *n* is the total number of sequences and *d*_*E*_(*i,j*) and *d*_*T*_(*i,j*) are respectively the estimated and true pairwise distances between sequences *i* and *j*. Because pairwise distances directly influence the accuracy of transmission clustering tools like HIV-TRACE, Mean Squared Error serves as a valuable indicator for the viability of any given MSA tool. We compute Mean Squared Error on pairwise distances computed directly from estimated MSAs under the Tamura-Nei 93 (TN93) model of sequence evolution (Tamura & Nei 1993) using the `tn93` component of HIV-TRACE (Pond *et al*. 2018), as well as from the pairwise distances along the inferred phylogenies.

### Mantel Correlation

We measure the correlation between the estimated and true pairwise distance matrices via the Mantel test. We include both Pearson and Spearman correlation, with 1 indicating perfect correlation (best), −1 indicating perfect anticorrelation, and 0 indicating no correlation.

### Robinson Foulds (RF) Distance

We use both normalized unweighted RF (URF) and weighted RF (WRF) distances to measure the topological difference between two unrooted trees (Foulds & Robinson 1981), with lower RF distances denoting higher accuracy.

### Sum of Pairs (SP) Score

SP Score measures the number of aligned pairs in the inferred MSA that are shared with the true MSA, normalized by the true MSA’s total number of homologies. Higher SP Score indicates higher accuracy. MSA SP Scores were computed using FastSP (Mirarab & Warnow 2011).

### Total Columns (TC) Score

TC Score measures the proportion of successfully-aligned columns out of the total number of aligned columns in the inferred MSA (Mirarab & Warnow 2011), which ranges from 1 (best) to 0 (worst). The TC Score of each MSA was computed using FastSP.

### Compression Factor

Compression Factor is defined as the number of columns in the inferred MSA divided by the number of columns in the true MSA. A perfectly-inferred MSA would have a compression factor of 1. MSA Compression Factors were computed using FastSP (Mirarab & Warnow 2011).

## Results

### Multiple Sequence Alignment

While Clustal Omega performs poorly on the HCV datasets and MUSCLE performs poorly on the Ebola datasets, MAFFT consistently performs the best across all three viruses in both runtime and accuracy (Fig. 1). Running MAFFT, MUSCLE, and Clustal Omega on the entire curated MSA of 2,322 complete HIV-1 genome sequences from LANL took 947 seconds, 20,343 seconds, and >86,400 seconds, respectively. All estimated MSAs of HIV-1 were slightly less accurate than those of HCV and Ebola with respect to all metrics aside from Mean Squared Error and Spearman Mantel correlation. Regardless, our results suggest that MAFFT is the best choice of the MSA tools that were tested in terms of both runtime and accuracy.

### Phylogenetic Inference

Despite the unweighted RF distance of Ebola phylogenies being markedly worse than the others, Ebola phylogenies appear to be comparable or better in accuracy according to all other metrics (Fig. 3). Additionally, using IQ-TREE under the GTR model performs similarly or better than using IQ-TREE with Model Finder Plus (MFP), and with a much lower runtime (Tab. 1), though it must be noted that the simulated MSAs were themselves evolved under the GTR model, which gives IQ-TREE GTR an inherent advantage over MFP. Phylogenies inferred from the HCV datasets have high Mantel correlations despite large Mean Squared Error (Fig. 3).

**Figure 3.**
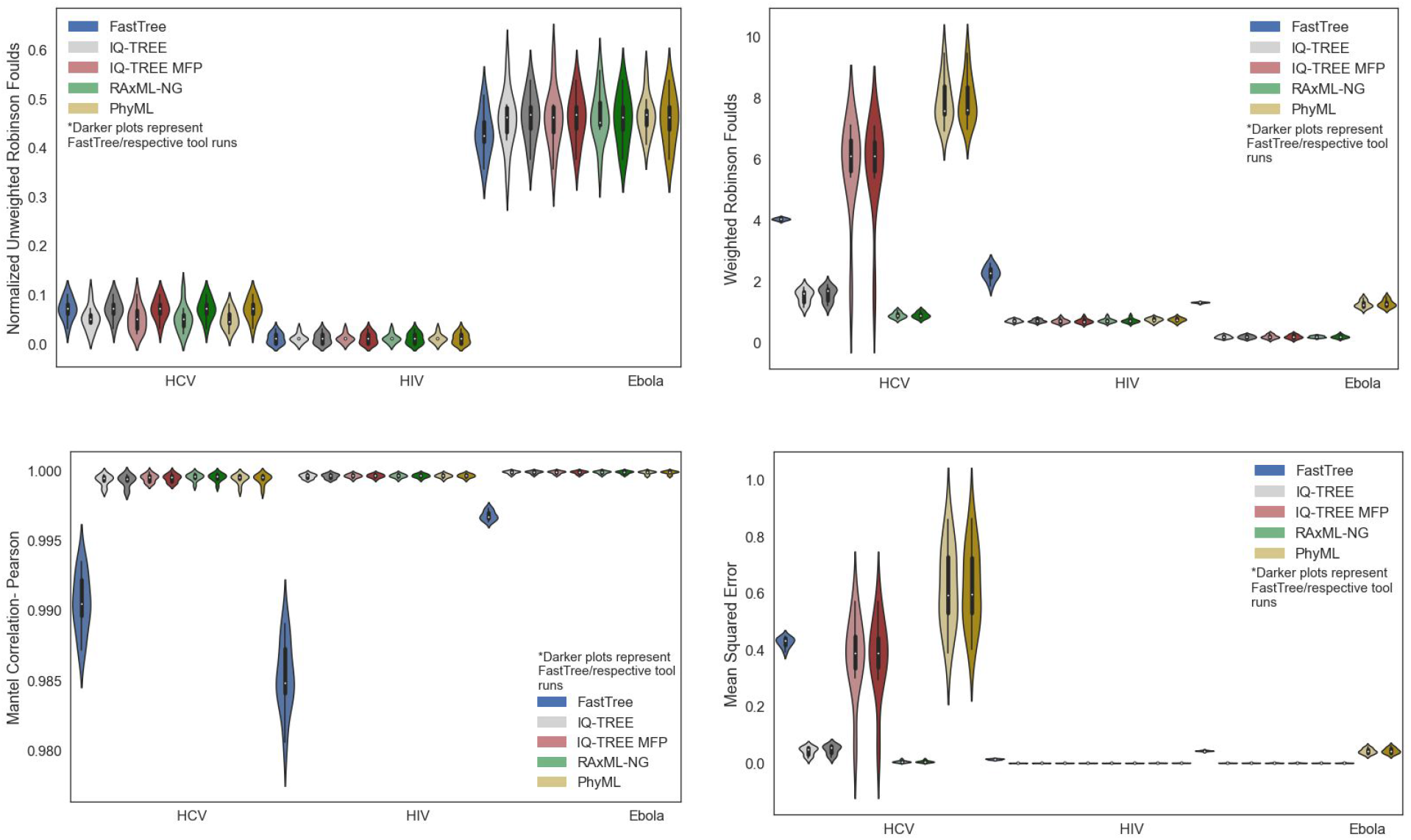
Metrics of phylogenetic inference accuracy for FastTree, IQ-TREE (GTR), IQ-TREE (MFP), RAxML-NG, and PhyML on 10 simulated replicate datasets of HIV, HCV, and Ebola. Phylogenies which result from optimizing branch lengths along FastTree topology are also included. Violin plots are shown for URF, WRF, Pearson Mantel Correlation, and Mean Squared Error.

As can be seen in Figure 3, FastTree 2 has notably worse accuracy than the other tools according to nearly all metrics across all three viruses, with the exception of Unweighted Robinson-Foulds, in which FastTree performs similarly. However, for each non-FastTree inference tool, the results of simply optimizing the branch lengths of a given FastTree topology has indistinguishable accuracy with respect to their counterparts in which tree search is performed by the same tool. Additionally, RAxML-NG performs comparably or better than other tools in terms of Weighted Robinson Foulds and Mean Squared Error.

### Combinations of MSA and PI

As expected, FastTree 2 consistently yields the lowest-accuracy phylogenies across all three viruses (Fig. 4). The other phylogenetic inference tools seem to perform similarly on HIV-1. Regarding Ebola, however, PhyML in particular struggles with Mean Squared Error and Weighted Robinson Foulds. As for HCV, IQ-TREE and RAxML-NG perform notably better than other tools. Model Finder Plus also seemed to misidentify the evolutionary model when used on HCV datasets, given that it performed markedly worse than IQ-TREE with GTR+I+R specified. Generally, the selection of MSA tool did not have a significant impact on the accuracy of the resulting phylogeny.

**Figure 4.**
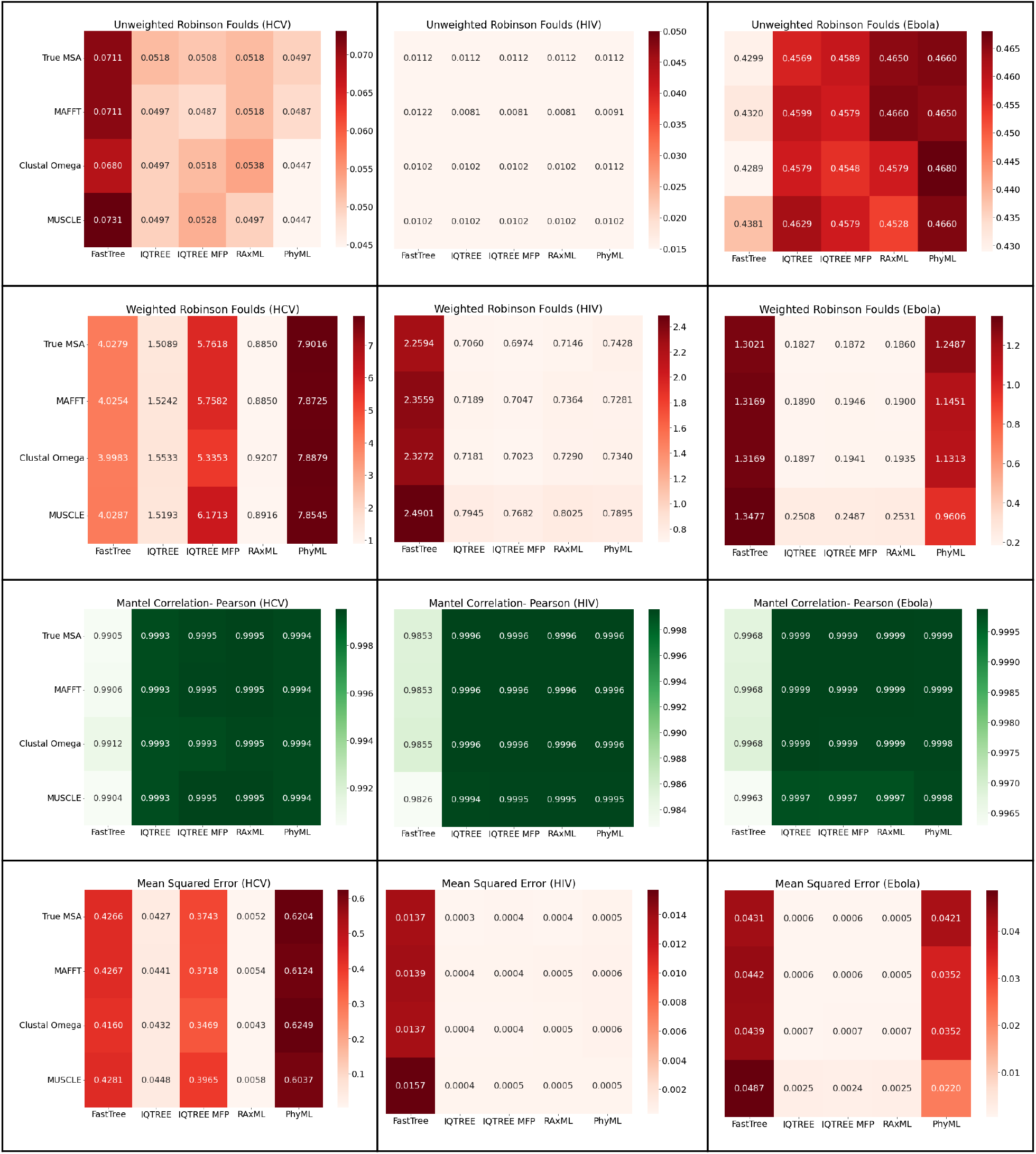
Heat maps comparing the accuracy of phylogenies inferred with FastTree, IQ-TREE (GTR), IQ-TREE (MFP), RAxML-NG, and PhyML from the MAFFT, Clustal Omega, MUSCLE, and true MSAs. Each value of Unweighted Robinson-Foulds (URF), Weighted Robinson-Foulds (WRF), Pearson Mantel Correlation, and Mean Squared Error shown is the average of 10 simulation replicates.

### Combinations of MSA and optimized FastTree topologies

IQ-TREE MFP struggles when inferring a phylogeny from the HCV datasets, having notably worse weighted RF distances and Mean Squared Error than IQ-TREE GTR (Fig. 5). In addition, the patristic distance matrices are highly correlated with the true matrix, even in instances where the other metrics may indicate poor performance.

**Figure 5.**
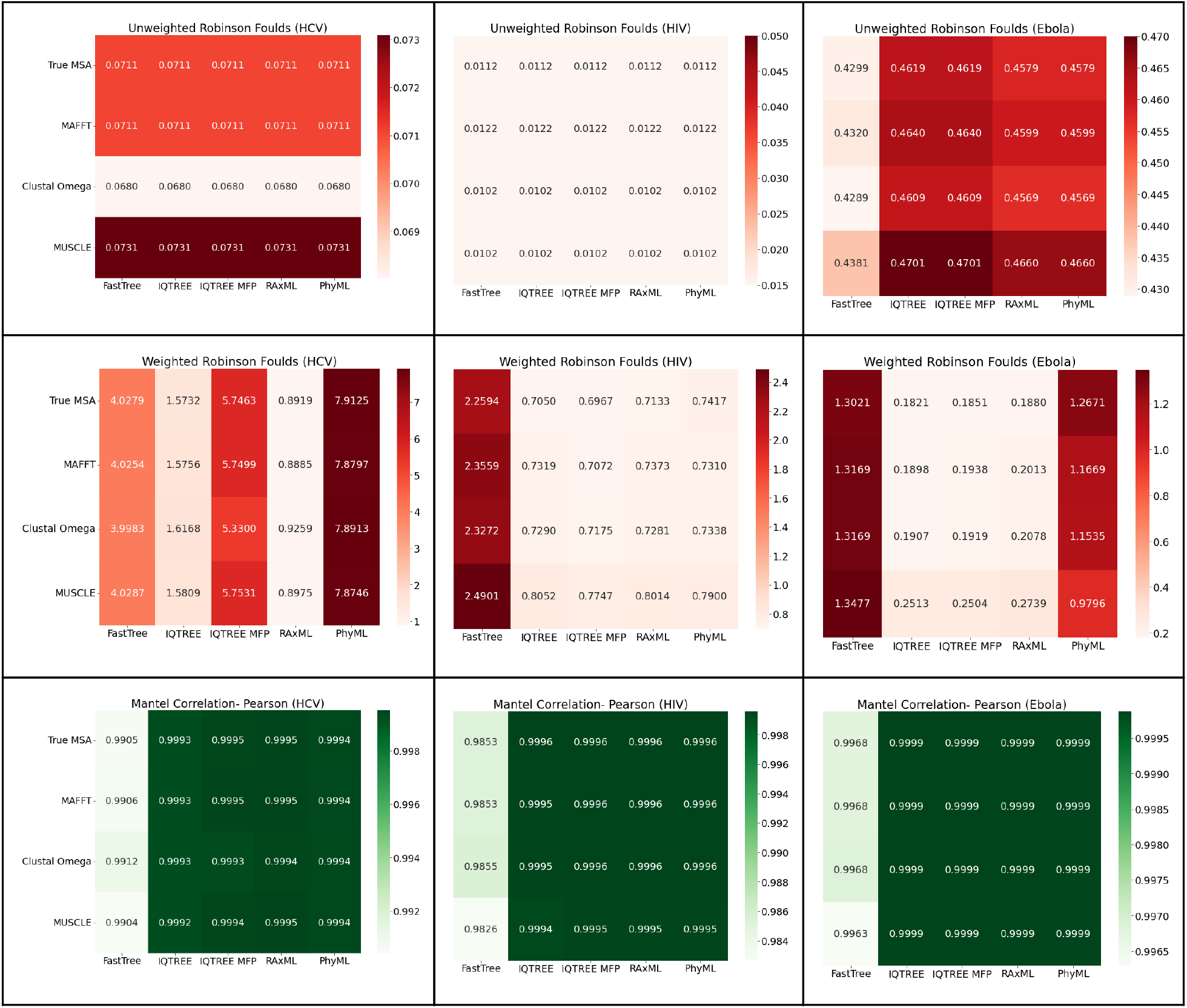
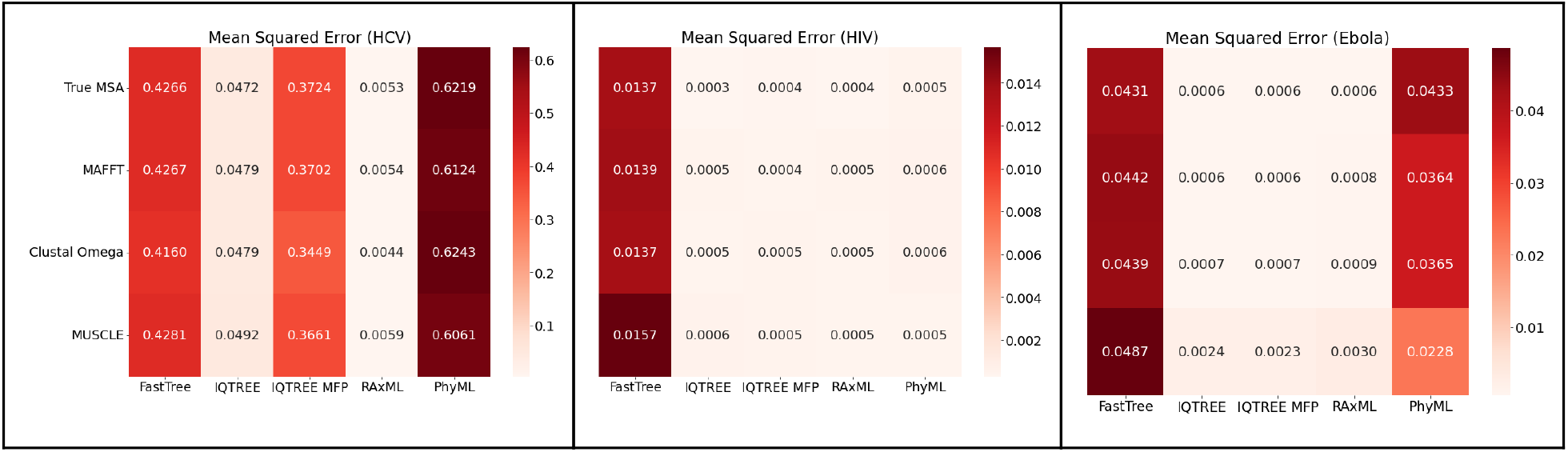
Heat maps comparing the accuracy of FastTree topologies inferred from the MAFFT, Clustal Omega, MUSCLE, and true multiple sequence alignments with branch lengths optimized by IQ-TREE (GTR), IQ-TREE (MFP), RAxML-NG, and PhyML. Each value of Unweighted Robinson-Foulds (URF), Weighted Robinson-Foulds (WRF), Pearson Mantel Correlation, and Mean Squared Error shown is the average of 10 simulation replicates.

Using the phylogenetic inference tools to optimize the branch lengths on a fixed topology significantly reduces the runtime across the board (Tab. 1), with the largest reduction in runtime being from RAxML-NG. IQ-TREE (GTR) and IQ-TREE MFP take notably longer just to optimize branch lengths, especially relative to their total runtime. With regard to tree search, FastTree 2 is several orders of magnitude faster than all the other inference tools.

## Discussion

Our results indicate that FastTree 2 consistently runs several orders of magnitude faster than other phylogenetic inference tools. With regard to accuracy, however, FastTree 2 performs worse than the other inference tools according to nearly all metrics across all three viruses, with the exception of Unweighted Robinson-Foulds. This suggests that FastTree 2 is able to infer a reasonably accurate tree topology, but that it poorly estimates branch lengths. Optimizing the branch lengths along a fixed topology inferred by FastTree 2 results in a phylogeny that is essentially identical to phylogenies inferred solely by a single tool, indicating that this workflow could considerably reduce the runtime of viral phylogenetic inference workflows with minimal loss in accuracy. It also suggests that each phylogenetic inference tool arrives at roughly equally-accurate topologies.

We also observed that phylogenies inferred for Ebola sequences have markedly worse unweighted RF distances than the phylogenies inferred for other viruses. However, the weighted RF distance for Ebola phylogenies actually measures lower. This is because a majority of the branch lengths in the true Ebola phylogeny were near zero, causing the weighting of each unique bipartition to be low, even if there were a significant number of unique bipartitions. The abundance of zero-length branches also intuitively made the phylogeny particularly difficult to infer, explaining the high unweighted RF distance.

Using IQ-TREE under the GTR model performed similarly to using IQ-TREE with Model Finder Plus when used on Ebola and HIV sequences, indicating that Model Finder Plus successfully classifies the evolutionary model in these instances. However, in HCV, IQ-TREE MFP correctly identified the model, transition rates, and proportion of invariable sites, but the relative rates in the model of rate heterogeneity were considerably inaccurate despite having the correct number of categories, resulting in a noticeable decline in accuracy. MFP also requires significantly more runtime, so unless the user has reason to believe that the GTR model is a poor fit for their specific virus of interest, the user is best off simply avoiding MFP.

Based on these results, our recommended workflow to obtain the highest-accuracy MSA and phylogeny with minimal runtime is to (1) use MAFFT to perform MSA, (2) use FastTree 2 under the GTR model with discrete gamma-distributed site-rate heterogeneity to quickly obtain a reasonable tree topology, and (3) use RAxML-NG to optimize branch lengths along the fixed FastTree 2 topology.

## Data Availability

All raw and processed data as well as scripts/tools utilized in this study can be found in the following GitHub repository: https://github.com/Cyoung02/Phylogenetic-Inference-Benchmarking

## Acknowledgements

This work has been supported by NSF grant NSF-2028040 to N.M., the Google Cloud Platform (GCP) Research Credits Program, and the UC San Diego Department of Biomedical Informatics Summer Training Program.

## Supplementary Materials

**Supplementary Fig 1.**
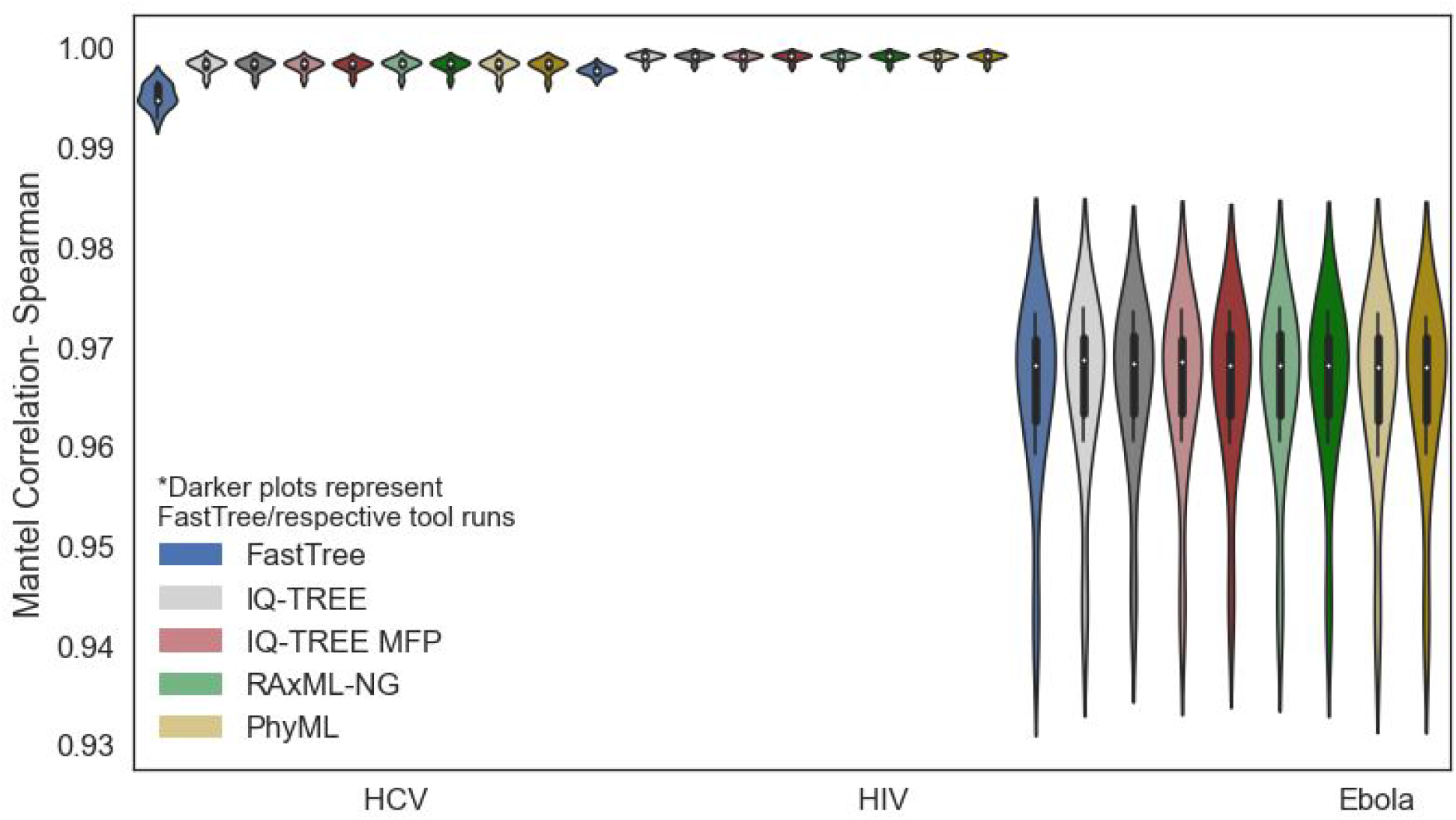
Violin plots of the Spearman Mantel Correlations for the patristic distance matrices of phylogenies inferred by FastTree, IQ-TREE, IQ-TREE (MFP), RAxML-NG, and PhyML from 10 simulated replicate datasets of HIV, HCV, and Ebola. Phylogenies which result from optimizing branch lengths along FastTree topology are also included.

**Supplementary Figure 2.**
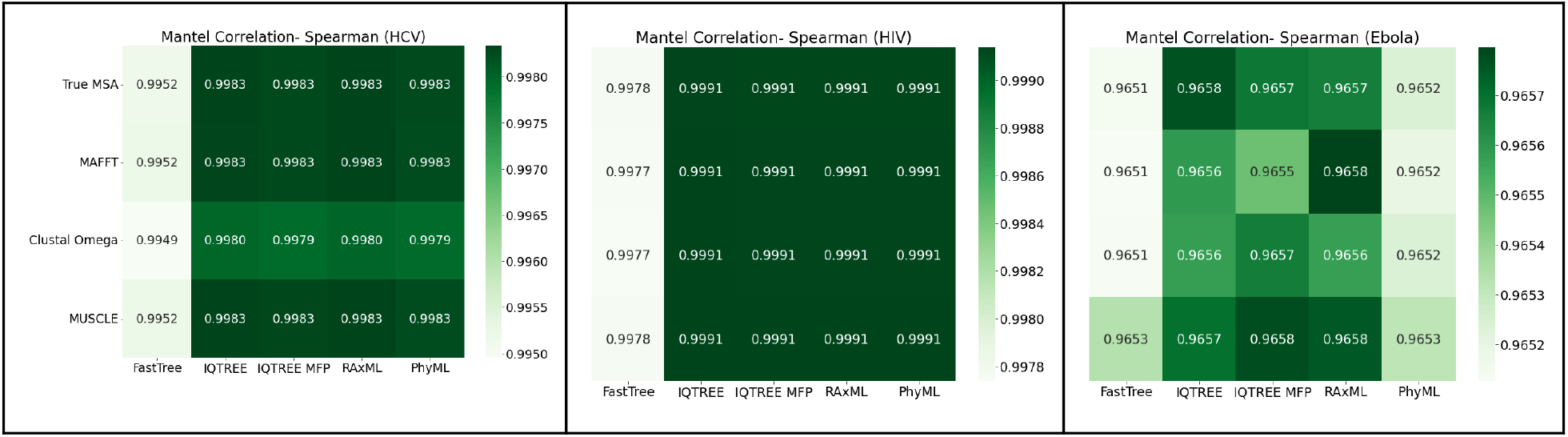
Heat maps comparing the average Spearman Mantel Correlations for patristic distance matrices of phylogenies inferred with FastTree, IQ-TREE, IQ-TREE (MFP), RAxML-NG, and PhyML from the MAFFT, Clustal Omega, MUSCLE, and true multiple sequence alignments. Values shown are the average of 10 replicate datasets for each virus.

**Supplementary Fig 3.**
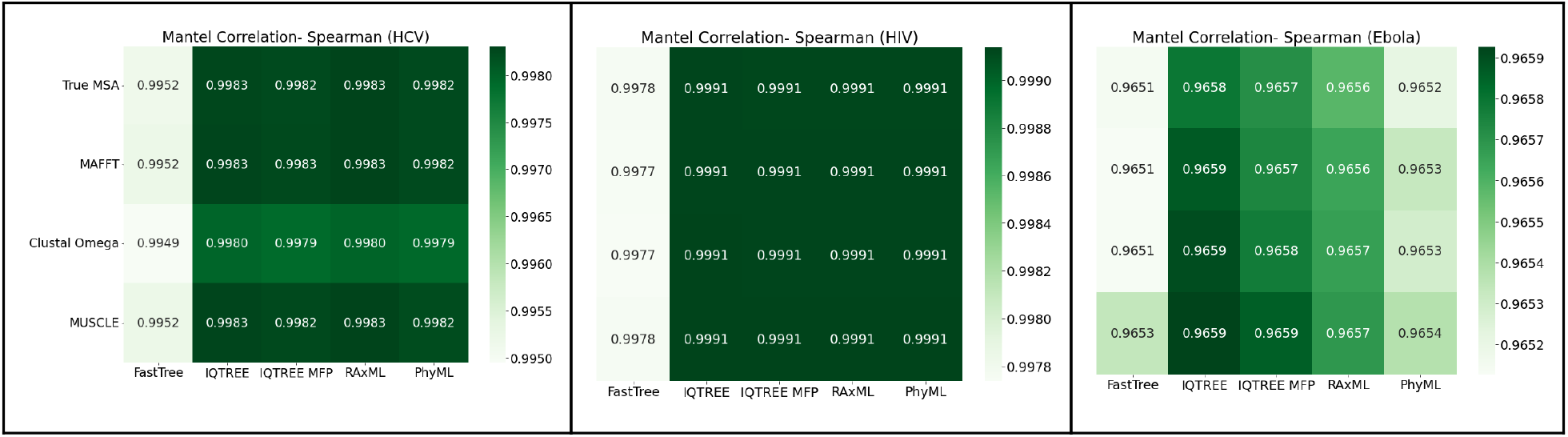
Heat maps comparing the Spearman Mantel Correlations for patristic distance matrices of FastTree topologies inferred from the MAFFT, Clustal Omega, MUSCLE, and true multiple sequence alignments with branch lengths optimized by IQ-TREE, IQ-TREE (MFP), RAxML-NG, and PhyML. Values shown are the average of 10 replicate datasets for each virus.

